# Effects of variable mutation rates and epistasis on the distribution of allele frequencies in humans

**DOI:** 10.1101/048421

**Authors:** Arbel Harpak, Anand Bhaskar, Jonathan K. Pritchard

**Author notes:** These authors contributed equally to this work (Arbel Harpak, Anand Bhaskar).

## Abstract

The site frequency spectrum (SFS) has long been used to study demographic history and natural selection. Here, we extend this summary by examining the SFS conditional on the alleles found at the same site in other species. We refer to this extension as the “phylogenetically-conditioned SFS” or cSFS. Using recent large-sample data from the Exome Aggregation Consortium (ExAC), combined with primate genome sequences, we find that human variants that occurred independently in closely related primate lineages are at higher frequencies in humans than variants with parallel substitutions in more distant primates. We show that this effect is largely due to sites with elevated mutation rates causing significant departures from the widely-used infinite sites mutation model. Our analysis also suggests substantial variation in mutation rates even among mutations involving the same nucleotide changes. We additionally find evidence for epistatic effects on the cSFS: namely, that parallel primate substitutions at nonsynonymous sites are more informative about constraint in humans when the parallel substitution occurs in a closely related species. In summary, we show that variable mutation rates and local sequence context are important determinants of the SFS in humans.

## Introduction

The distribution of allele frequencies across segregating sites, commonly referred to as the Site Frequency Spectrum (SFS), is a central focus of population genetics research as it can reflect a wide range of evolutionary processes, including demographic history as well as positive and purifying selection [1–8]. Until recently, the SFS was usually measured in samples of tens or hundreds of people, but advances in sequencing technology have enabled the collection of sequence data at much larger scales [9–14]. Notably, the Exome Aggregation Consortium (ExAC) recently released high quality, exome-wide allele counts for over 60,000 people [12].

Large sample sizes are valuable because they make it possible to detect many more segregating sites, and to estimate the frequencies of rare variants. For example, the recent dramatic expansion of human populations leaves little signal in the SFS in small samples [15], but is readily detected in large samples, where there is a huge excess of low frequency variants compared to model-predictions without growth [13,14,16,17]. Similarly, large samples enable the detection of deleterious variants that are held at very low frequencies by purifying selection [18–22].

In this paper, we extend the SFS by considering the SFS *conditional* on the observed alleles at a given site in other species (specifically, other primates in our analysis). Our original motivation was that this could allow us to measure the effects of sequence context on the selective constraint of missense variants. In general, sites with strong levels of average constraint across mammals tend to be less polymorphic within humans [16,23,24], but to the best of our knowledge, there has not been extensive consideration of the joint distribution of the substitutions across other lineages and the human SFS. In particular, we hypothesized that if an identical substitution has occurred independently in a closely related species—e.g., in a great ape—then this is strong evidence that the same variant is unlikely to be deleterious in humans. However, an identical substitution in a more distantly related species may be much less informative, as substitutions at other positions within the same gene may change the set of preferred alleles due to epistatic interactions [25–31] (Figure 1). For example, it has been shown that, in a handful of cases, likely disease-causing variants in humans are actually wildtype alleles in mouse, presumably rendered harmless by parallel substitutions at interacting positions [26].

**Figure 1.**
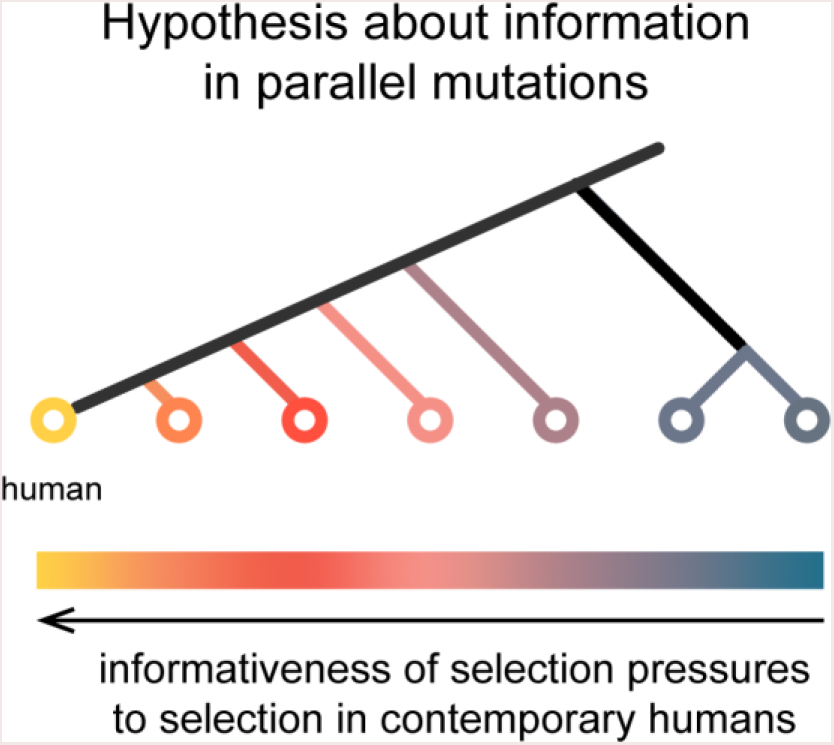
Hypothesis about information in parallel mutations. If an identical substitution occurred independently in a closely related species, then the variant is unlikely to be deleterious in humans. An identical substitution in a more distantly related species may be less informative because sequence divergence at interacting sites may change the set of preferred alleles, and hence the selective constraint at the site.

As we show below, the human SFS varies greatly depending on patterns of substitutions in other species. In part, this does appear to be due to the accumulation of epistatic effects on more distant lineages; however a more important factor is mutation rate variation across sites. Under the widely-used infinite sites model, the SFS is independent of mutation rate; but in the ExAC dataset we observe a clear breakdown of this model. Mutation rates are known to vary across sites due to a variety of different mechanisms, leading to differences between CpGs, transitions and transversions, as well as additional effects that correlate with broader sequence context, replication timing, transcription, recombination rate and chromatin environment [32–39]. We show here that mutation rates are much more variable than generally appreciated, and that rates at some sites are high enough to generate substantial deviations from infinite sites predictions. The main ExAC paper [12] also recently reported that the SFS varies substantially across mutation types, and also noted that this implies departures from the infinite sites model, especially for CpGs.

In summary, our results suggest more variation in mutation rates across sites than is generally appreciated, and further that the infinite sites model provides a poor fit for population genetic analyses in large modern data sets. We also show a significant, albeit smaller, role for epistatic effects in shaping the cSFS.

## Results

To investigate the properties of the human cSFS, we combined exome sequence data from 60,706 humans from ExAC version 0.2 [12,40] and orthologous reference alleles for 6 nonhuman primate species from the UCSC genome browser [41]. After applying several filters (see **Materials and methods** for details) we were left with 6,002,065 single nucleotide polymorphisms (SNPs) for which we had orthologous data in at least one nonhuman species.

We examined how the human SFS changes as we condition on various divergence patterns observed in primates. There are many possible ways to condition on variation across the nonhuman primates. We focus here on sites that are variable in human only (denoted *human-private*), as well as sites where exactly one other species carries the human minor allele (and all others match the human major allele); see Figure S1 for an alternative conditioning based on the most closely related species carrying the human minor allele. Throughout, we assume that the observation of the human minor allele as the reference allele in another primate implies that the mutation arose independently and fixed in that primate. This assumption may be violated for a small fraction of SNPs when comparing human to our closest relatives (notably, chimpanzee and gorilla [42]), but the overall patterns that we report here are maintained when we consider more distant species for which shared ancestral polymorphism is unlikely (see **Materials and Methods** and **Supplementary Material** for further discussion). The SFS presented here, unless otherwise stated, are constructed using minor allele frequencies.

Henceforth, we will use the term *substituted species* to refer to the single species in which the human minor allele is observed, and the corresponding *species cSFS* to refer to the human SFS conditional on a substituted-species divergence pattern. For example, “substituted-orangutan” refers to human variants for which the human minor allele is observed in orangutan, and the human major allele is observed in all other primates; “orangutan cSFS” will refer to the human SFS at these sites (Figure 2A). There were 5,286,937 human-private sites in the data set, and the number of substituted-species sites ranged from 22,209 (substituted-chimpanzee) to 66,254 (substituted-gibbon).

**Figure 2.**
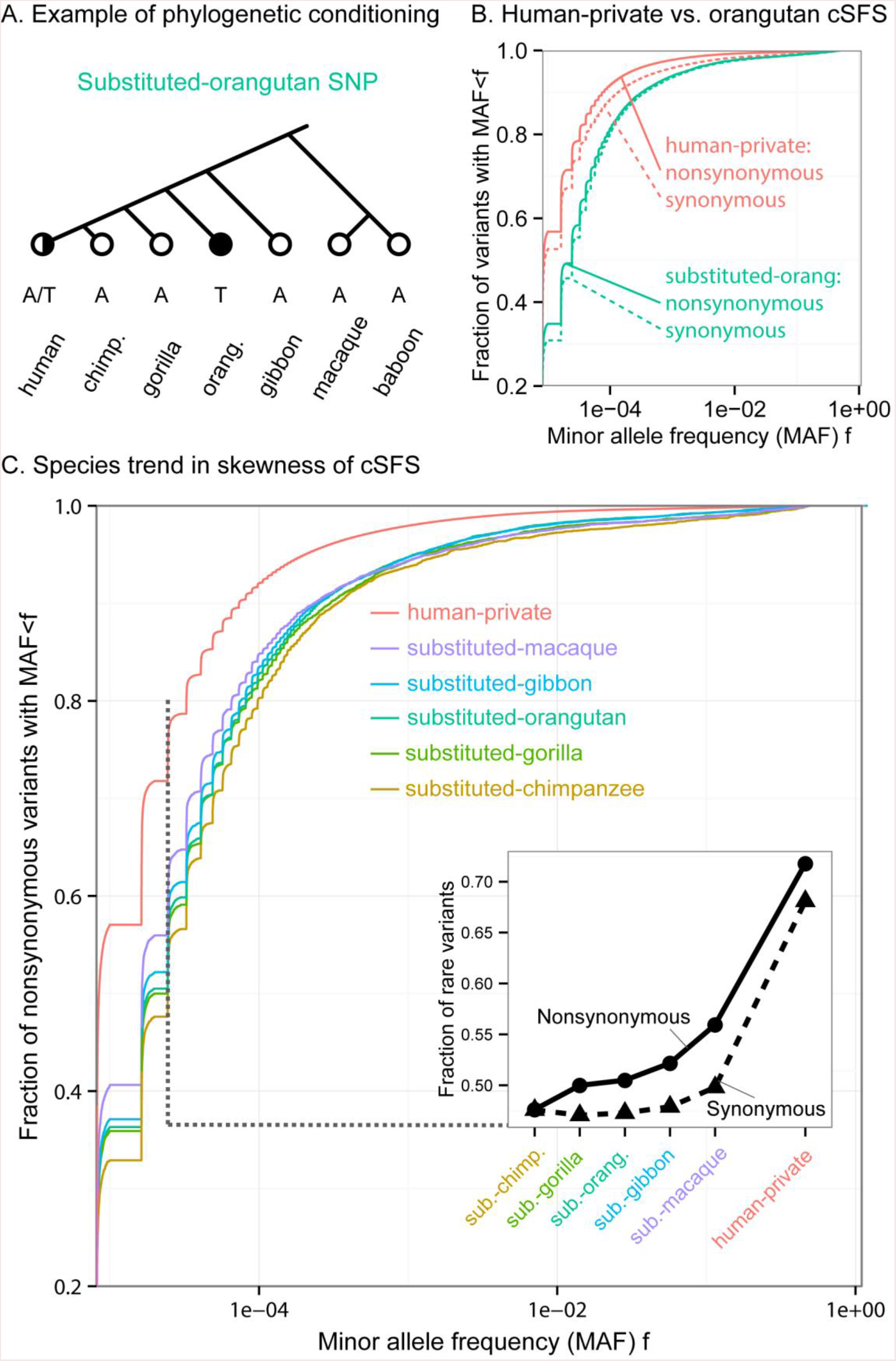
The human SFS conditioned on primate substitution patterns. **(A)** An example of the phylogenetic conditioning that defines what we denote as “substituted-orangutan” sites. **(B)** The cumulative distribution functions (CDF) of orangutan cSFS (i.e. the SFS of substituted-orangutan sites), and the SFS of phylogenetically conserved sites. The cSFS are more skewed towards common variants than the SFS of conserved sites. These skews are much more pronounced than in the comparison of synonymous and nonsynonymous sites. **(C)** The more closely related the substituted-species, the higher the skew of the cSFS towards common variants (only nonsynonymous mutations shown). The inset shows the rare variants slice of the CDF for each species, for both synonymous and nonsynonymous variants.

Figure 2B shows a comparison of the human-private cSFS and the orangutan cSFS for nonsynonymous and synonymous sites, respectively. Within each cSFS class, the nonsynonymous spectrum has more rare variants than the synonymous spectrum, as expected given that nonsynonymous variants are more likely to have deleterious effects. Secondly, if we compare the human-private versus orangutan cSFS at nonsynonymous sites, we see more rare variants in the human-private set. Again, this matches expectations, as the presence of a parallel substitution in orangutan implies that a substitution at this position is tolerated.

However, we were surprised to see that substituted-orangutan synonymous sites also segregate at much higher frequencies than both synonymous and nonsynonymous human-private sites. Taken at face value, this would seem to imply that a large fraction of synonymous sites are functionally constrained. While it is known that some synonymous sites play roles in functions such as splicing [43,44], it is generally believed that most synonymous variants in mammals are effectively neutral. We were thus curious to understand whether this result is primarily driven by a surprising degree of constraint at synonymous sites, or by some other factors.

Looking more broadly across the primates, we observed a clear trend of cSFS across substituted species (Figure 2C): the more closely related the substituted species, the greater the skew towards high frequency variants. This trend is most easily noticeable in the fraction of rare variants (defined here, arbitrarily, as singletons and doubletons; Figure 2C, inset). In the following sections, we try to understand the factors driving these observations.

### Effect of mutation rate variation on the human SFS

In this section we consider whether mutation rate variation may contribute to the observed trend across cSFS. Under the standard infinite sites assumption, the SFS is independent of mutation rate. However, we conjectured that in the very large sample size of ExAC, infinite sites may no longer be a good model for the data [45].

To examine this, we stratified the human SFS by mononucleotide mutation types (as well as the dinucleotide mutation type CpG->TpG), for which there are well-characterized differences in mutation rates. For this analysis we focused on intronic sites, to reduce potential effects of selective constraint. We found that the different mutation types have significantly distinct spectra. The fraction of rare variants among CpG->TpG mutations (36%) was roughly half that of non-CpG transitions (71%, see Figure 3A). Similarly, non-CpG transitions have higher mutation rates than transversions and indeed, the SFS for transitions is also skewed towards higher frequencies than transversions (Figure 3B). Overall, the fraction of rare variants in the subsample of Europeans was significantly negatively correlated with germline mutation rates estimated from the deCode project dataset [46] (weighted linear regression *p* = 4.9×10^−6^, see Figure 4A and **Supplementary Material**).

**Figure 3.**
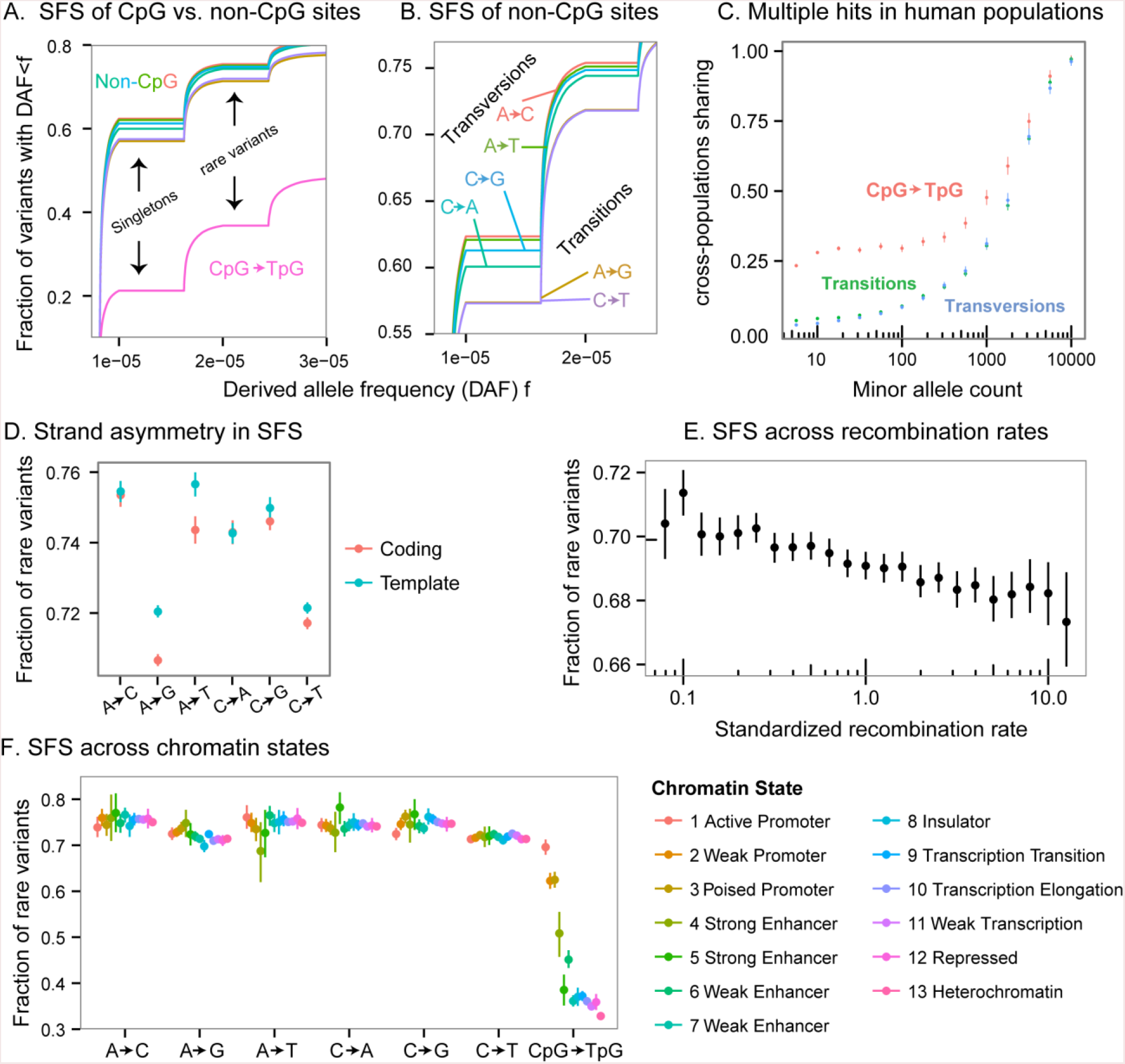
Rare variant frequencies vary dramatically by mutation type. All panels show the SFS of derived alleles constructed from intronic sites. The notation of mutation types refers to mutations on either strand (e.g., A->C indicates an A to C change on either strand). **(A)** SFS stratified by non-CpG mononucleotide mutation types and CpG transitions, represented by different curves. The fraction of rare variants in CpG transitions is nearly half that of other mutations. **(B)** Focusing on non-CpG mutations, transitions have an SFS significantly skewed towards common variants compared with transversions. **(C)** Sharing of polymorphisms between East Asians and Europeans. The excess sharing of CpG polymorphisms at low frequencies is suggestive of multiple occurrences of the mutations. x-axis values are binned on a logarithmic scale. **(D)** Stratification to coding and template strands revealed differences between the two for some mutation types, suggesting transcription-associated mutational mechanisms also affect the SFS. CpG mutations excluded from the analysis in this panel. **(E)** Recombination rates are positively correlated with the fraction of rare variants; this could be due to a correlation between recombination rates and mutation rates. x-axis values are standardized to the genomewide mean, and are binned on a logarithmic scale. **(F)** SFS across chromatin states. Chromatin states in H1 human embryonic stem cells were inferred by ChromHMM. The chromatin state exhibits substantial association with the fraction of rare variants in CpG mutations, and modest association in other mononucleotide mutation types. In panels D,E and F: Points show means; lines show 95% confidence interval computed with nonparametric bootstrap.

**Figure 4.**
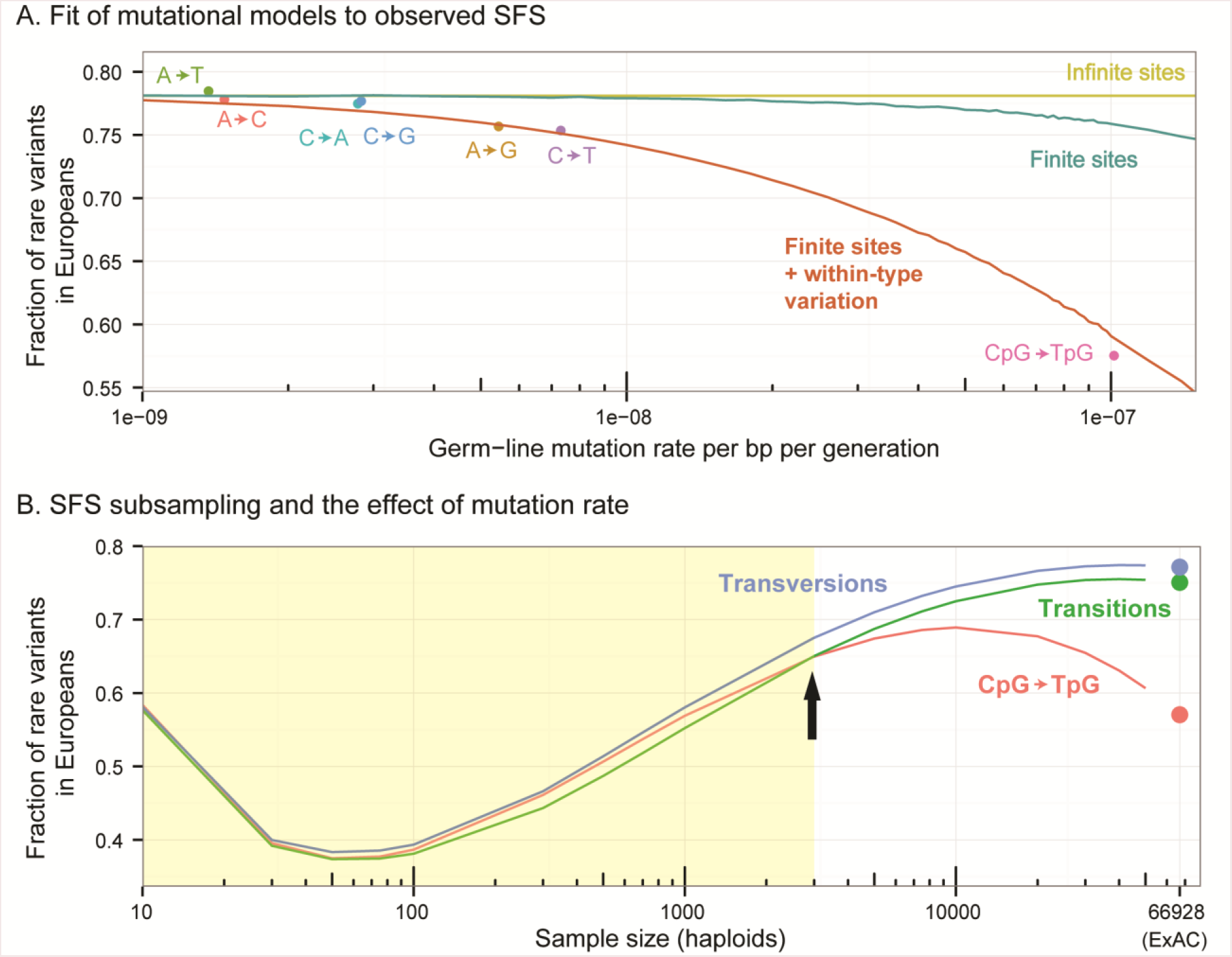
All panels exhibit the unfolded SFS (i.e., constructed using the derived alleles) of intronic sites. **(A)** Fit of mutational models to observed SFS. The x-axis shows previously estimated de-novo germ-line mutation rates [46]. These data illustrate that the fraction of rare variants is strongly negatively correlated with germ-line mutation rates. Lines show expectations under various mutational models: yellow–infinite sites model (SFS independent of mutation rate); teal–Jukes Cantor finite-sites model; red–Jukes-Cantor model with within-mutation-type variation (i.e., variation beyond mutation rate heterogeneity due to the type of mutation in sequence). **(B**) SFS subsampling and the effect of mutation rate. Dots show the fraction of rare variants in the full sample SFS of the European population in ExAC. Lines show the expected fraction of rare variants after subsampling to smaller numbers of individuals. In large samples, the SFS of CpG and non-CpG sites are very different. In smaller samples, these differences shrink. In the shaded region, the trend across mutation types is changed (the inflection point is indicated by an arrow); with these sample sizes, CpG transitions exhibit more rare variation than non-CpG transitions.

If multiple hits are prevalent within the ExAC sample, some of them should occur in different subpopulations. Higher mutation rates should then lead to excess sharing of low frequency variants among subpopulations. To verify the occurrence of recurrent mutations, we examined the sharing between the European and East Asian ExAC subsamples. Indeed, at low frequencies, non-CpG transitions exhibited a higher sharing rate than transversions, and CpG transitions exhibited much higher sharing rate than non-CpG sites (For example, for sites with minor allele count of 10, we get a t-test *p* < 2.2 · 10^−16^ for both comparisons; see Figure 3C, and a similar analysis performed in Figure 2d in the main ExAC paper [12]).

As an additional test of whether mutation rate affects the fraction of rare variants, we turned to sites in transcribed regions. It is known that in such regions, A->G and A->T mutations occur at higher rates on the template (non-coding) strand than on the non-template (coding) strand, due to the effects of transcription-coupled repair or other transcription-associated mutational asymmetries [47–49]. Indeed, as predicted from these rate asymmetries, we observed a 1% difference between the template and the coding strands in the fraction of rare variants in introns (t-test *p* < 2.2 · 10^−16^ for A−> G, *p* = 6.0×10^−7^ for A->T). C->T mutations also exhibit a small but significant difference (t-test *p* = 3.0×10^−4^) between the strands, even though, to our knowledge, no previous work has observed a rate asymmetry for C->T mutations (Figure 3D).

Similarly, we hypothesized that the SFS at CpG sites might also depend on chromatin environment (Figure 3F). Specifically, CpG sites experience high mutation rates only when they are methylated [50–53]. We thus examined the effect of chromatin states in H1 human embryonic stem cell lines, inferred by ChromHMM [54] (as a proxy for germline chromatin states) on the SFS across different mutation types. Methylation levels are expected to be low in active regions including promoters and enhancers and high in repressed regions such as heterochromatin. Indeed, we find highly significant differences in the SFS at CpGs (see **Supplementary Material** for details), consistent with this expectation: i.e., fewer rare variants in heterochromatin, where methylation levels are high. In contrast, the other mononucleotide mutations showed only modest variation across chromatin states.

Finally, we found that recombination rate is also negatively correlated with the fraction of rare variants (Pearson correlation *p* < 2.2 · 10^−16^, and see Figure 3E and Figure S5). This is consistent with the postulated positive correlation between recombination and mutation rates [55,56]. However, linked selection—which is expected to be more pervasive in regions of low recombination—could also contribute to this trend [57–59]. Overall, the SFS variation patterns across chromatin states, recombination rates, and strands, underscores that heterogeneity in mutation rates does exist within mutation types, and that it has a substantial effect on the SFS.

These observations on mutation rate variation led us to conclude that the infinite-sites model provides a poor fit for these large-sample human polymorphism data. We therefore investigated finite-sites mutational models. Below, we describe the fit of various mutational models while using previously-inferred population genetic models of European demography. In particular, we eventually used a modified version of the demography inferred by Nelson et al. [14] (see **Materials and Methods** for the other demographic models considered). The assumed demography provides a good fit for the SFS of sites with the lowest mutation rates.

We asked how well different finite-sites models account for the observed relationship between de-novo mutation rates and the SFS. First, we considered the Jukes-Cantor model, which uses a 4 × 4 uniform mutation transition matrix [60]. But we were surprised to find that this finite sites model barely improved the fit to the SFS across the range of estimated mutation rates (Figure 4A). In our simulations, the probability of obtaining more than one mutation on the genealogy of a segregating site is low enough that the finite sites SFS is similar to the infinite sites SFS, even at the relatively high mutation rate estimated for CpGs.

We hypothesized that we might achieve a better fit if some sites have higher intrinsic mutation rates than the mean for the particular nucleotide change at that site; this notion has received increasing support in the recent decade from both evolutionary and family-based studies of human mutation rates [32–35,37,61–63]. We therefore augmented the Jukes-Cantor model by incorporating additional variation in mutation rates across sites belonging to each mutation type (see **Materials and Methods**). The augmented Jukes-Cantor model with within-mutation-type variation fitted the data well, including the large difference in SFS between CpG and non-CpG sites (Figure 4A). The augmented model suggests that 3% of mutations within a mutation type have a mutation rate of over 5 times the mean rate for that type. This estimate is close to the level of mutation rate variation inferred by Hodgkinson and Eyre-Walker [64].

It is natural to wonder what effect recurrent mutations may have in smaller samples. Small samples have the disadvantages of increased noise and limited temporal resolution of analysis. For example, in demographic inference, larger samples are essential for detecting the signal of recent rapid growth of the human population [17,65,66]. Interestingly, we found that samples much smaller than ExAC may also create an unappreciated bias, as we describe next.

We examined the effect of subsampling the SFS of the European ExAC sample to a smaller number of individuals (see **Supplementary material**). SFS differences between non-CpG transitions and transversions remained roughly the same, even with a sample of a few hundred people. Conversely, the difference between CpG and non-CpG sites changed dramatically for smaller samples. For samples smaller than 1500 people, there appears to be more rare variation in CpG than non-CpG transitions (Figure 4B, Figure S4). This finding exemplifies that if one category of sites has substantially more rare variation in the population than a second category, the sample SFS may actually exhibit more rare variation in the second category. Therefore, a comparison of the amount of rare variation across categories of sites may yield different orderings, depending on the sample size.

Finally, we returned to the species trend across cSFS that we described earlier (Figure 2C). Given the previous observations on SFS differences between mutation types, we asked whether the trend across substituted-species cSFS we described earlier (Figure 2C) could be explained by differing compositions of the various mutation types. Indeed, most of this trend is due to the fact that CpG transitions make up a higher fraction of sites for more closely related substituted species (Spearman *ρ* = −0.9, *p* = 0.08, and see Figure S2). Since CpG transitions are depleted of rare variants, this results in the cSFS skewness trend. Namely, the fraction of rare variants is strongly negatively correlated with the fraction of CpG transitions across substituted-species (Pearson *r* = −0.997, *p* = 9.7 10^−6^ for nonsynonymous mutations; *r* = −0.999, *p* = 9.9×10^−7^ for synonymous mutations, see Figure 5C).

**Fig 5.**
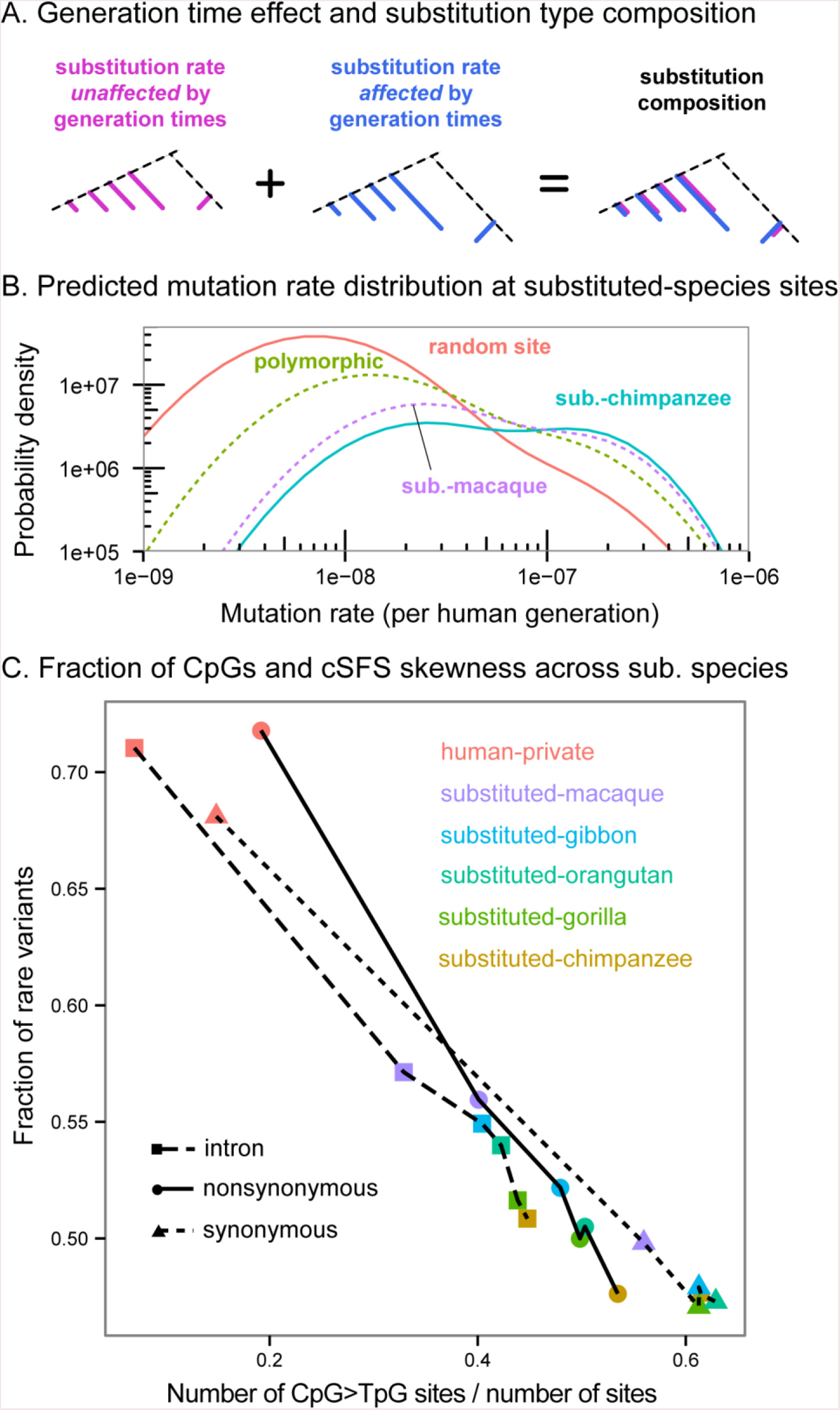
**(A)** Some mutation types accumulate in a roughly constant yearly rate across different primate lineages. For these mutation types the expected number of substitutions on an evolutionary branch is proportional to the branch length in years (pink). The yearly rates of other substitution types (blue) depends on various life-history traits like generation times (“generation time effect”). As a result, the composition of substitution types in a lineage depends on lineage-specific traits like generation times; this is illustrated by the blue to pink ratio, which differs across lineages. **(B)** Model-based expectations for the distribution of mutation rates at substituted-species sites. These results were computed using a theoretical model and a set of realistic parameters. At substituted species sites, we expect a distribution skewed towards higher mutation rates compared to random sites, or to random polymorphic sites. In addition, the distribution of mutation rates is skewed towards higher mutation rates for substituted-species with longer generation times; for the primates we considered in this work, this would imply higher mutation rates for more closely-related substituted-species. **(C)** CpG transitions enrichment is a strong predictor of cSFS skewness in real data.

Why is the fraction of CpG transitions negatively correlated with the relatedness of the substituted-species to humans? Below, we suggest how this could be explained through the mutational mechanism of CpG transitions, which leads to different substitution dynamics on evolutionary timescales than the dynamics at non-CpG sites.

Substituted-species sites likely experienced two independent mutations at the site during primate evolution, and are therefore enriched for hypermutable sites [61,62,64]. A simple model that we develop in the **Supplementary Material** supports this intuition. In this model, we initially assumed a “uniform molecular clock” regime in which substitutions accumulate at the same yearly rate across the primate phylogeny. Under this assumption, differences in the distribution of mutation rates between substituted-species categories should be vanishingly small (Figure S6).

However, recent work [67–70] has demonstrated that while the “uniform clock” assumption is valid for some mutation types—importantly, CpG transitions—the yearly substitution rates of other mutation types depend heavily on life-history traits such as generation time [69,71,72], and thus vary extensively across primates. Notably, Moorjani et al. have pointed out that this difference leads to variable mutational spectra across primates [67]. We therefore augmented our model by including two mutation categories: mutation types that follow a “uniform clock”, and mutation types with rates that depend on generation times (**Supplementary Material,** and see Figure 5A). The model predicts an enrichment of uniform-clock mutations for substituted-primates with longer generation times. Notably, this translates into a prediction of an enrichment of uniform-clock mutations—like CpG transitions—in substituted species more closely-related to humans (with the exception of orangutan, which is thought to have the longest generation time among the primates considered, although it was only estimated in females [73]). Examining the expected distributions of mutation rates in substituted-species sites, this enrichment leads to a skew towards higher mutation rates for more closely-related substituted-species (Figure 5B).

Overall, this model provides an explanation by which mutational mechanisms underlie the observed correlation between the relatedness of the substituted species and the skew of its cSFS towards common variants. We next asked whether additional causes beyond mutation rate variation might also contribute to the species trend across cSFS.

### Effects of epistasis on the human SFS

A second process that could contribute to the observed pattern of cSFS differences across substituted-species is fitness epistasis [25–27,31]. It is well-known that sites that are functionally important in humans tend to be relatively conserved across the mammals [74]. However, this is neither a necessary nor a sufficient condition for predicting functional sites in humans, and there are some counter-intuitive examples of disease-causing mutations in humans that are annotated as the reference allele in mouse [26]. It is presumed that such cases may be explained by parallel changes at other interacting amino acids that alter the structural context of the relevant site in mouse.

We thus hypothesized that the observation of a human variant in a closely related species provides suggestive evidence that the allele may be benign for humans as well. This evidence is expected to be stronger the more closely related the other species is to humans, because there would have been less time to accumulate additional epistatic interactions. An effect of this type could contribute to the trends observed in Figure 2C. In this section we test for evidence of epistatic effects.

To this end, we used a logistic regression model (see **Materials and Methods**). We first examined whether the probability of the variant being rare is associated with the relatedness of the substituted species. A model that included only the relatedness of the substituted species showed a perfectly correlated ordering of the two (Figure 6A, CpG transitions were excluded from this analysis). We then turned to examine whether this correlation persists after controlling for mutational composition differences between substituted-species categories. We controlled for the effect of mononucleotide mutation types on the probability of the variant being rare (Figure 6B). We then further refined the mononucleotide mutation types by using their two flanking nucleotides, and estimated another model with these finer mutation type categories (Figure 6C). The trend persisted even after controlling for mutation type, most noticeably for nonsynonymous sites, which likely involve the strongest purifying selection pressures (Spearman *ρ* = 1, *p* = 0.016 for the ordering of substituted-species coefficients for both models). We repeated the analysis while including CpG transitions, and found that the perfect correlation persisted (Spearman *ρ* = 1, p=0.016 for both models; Figure S12).

**Figure 6.**
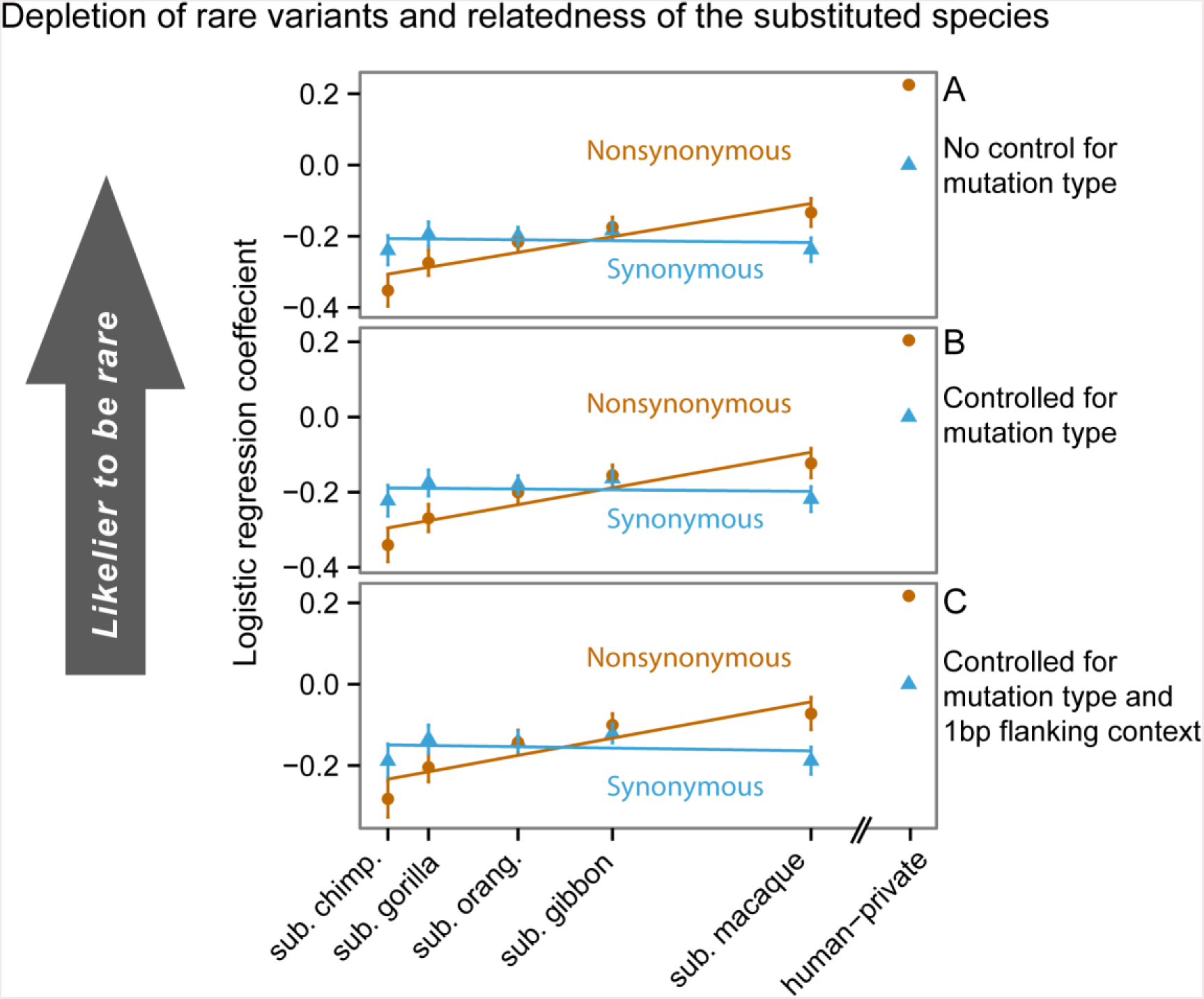
Depletion of rare variants is correlated with relatedness to substituted species. The figure shows logistic regression coefficient estimates with their corresponding standard errors. Substituted-species labels are spaced by their split times from humans. The lines are the least-squares line fitting the coefficients to the split times. **(A)** Estimates from a simple logistic regression to the substituted species. The trend is partly due to mutational composition differences between substituted-species categories. To test whether the trend is driven solely by mutational rate differences, we estimate coefficients in a model including the variation explained by **(B)** mononucleotide mutation type, and **(C)** combinations of focal mononucleotide mutations and upstream and downstream nucleotides. Even after controlling for mutational composition with these models, a significant trend persists for nonsynonymous variants.

One explanation for the residual trend observed in Figures 6B,C is that more-related species have, on average, more similar context on which the mutation occurs. We can interpret this residual trend as support for the following hypothesis: when the sequence context of the substituted species is similar to that of humans, the fixation of the human-minor allele in the substituted species suggests that the mutation is benign for humans. As sequence context diverges, epistatic effects may come into play and change the selective effect of the mutation [28,75,76]. In the **Supplementary Material**, we investigate the effect of sequence context divergence more directly.

## Discussion

Our analysis showed a significant correlation between the probability that a variant is rare in humans and the relatedness of another species in which the same mutation occurred. This trend was largely driven by mutation rate variation, which we have observed to be a primary determinant of the human SFS.

The large effect that mutation rate variation has on the human SFS could have a major impact on any future work involving human polymorphism datasets with large sample sizes. For example, most demographic inference algorithms that use the SFS as a summary statistic [e.g. 6,66,77] rely on the infinite-sites model, which is evidently not a valid assumption for large samples. Adjusting demographic inference schemes to include the effects of recurrent mutations on the SFS (for examples of recent efforts towards this goal, see [78–82]) has the potential to significantly improve inference accuracy.

We have also seen that the trend across cSFS persisted even after tri-nucleotide mutational composition was taken into account. This remaining correlation is consistent with an effect of sequence context epistasis on the fate of mutations.

Substitutions in other lineages have proven to be highly informative for understanding deleterious effects in the contemporary human genome; among numerous features that have been considered, the strongest predictors of the pathogenicity of a mutation are species divergence features [83–86]. Nevertheless, methods used to predict the deleteriousness of a mutation at a site typically rely on a single summary of how variable a site is across the phylogenetic tree. Our analysis suggests that epistatic effects can bias the inferred deleteriousness of the mutation, and that the location of a mutation on the evolutionary tree is informative of how deleterious the same mutation is for humans. It is our hope that the integration of divergence patterns and sequence context into methods that predict the fitness or health effects of human mutations could increase accuracy and predictive power.

## Materials and Methods

### Data

For polymorphism data, we downloaded single nucleotide polymorphism (SNP) data from version 0.2 of the Exome Aggregation Consortium database [40]. This database is a standardized aggregation of several exome sequencing studies amounting to a sample size of over 60,700 individuals and approximately 8 million SNPs. For each SNP we extracted upstream and downstream 30 nucleotides in the coding sequence of the human reference genome hg19 build. For simplicity we excluded sites that are tri-allelic (6.5% of all SNPs) or quad-allelic (0.2% of all SNPs).

For divergence data, we used the following reference genome builds downloaded from the UCSC genome browser [41]: chimpanzee (panTro4), gorilla (gorGor3), orangutan (ponAbe2), gibbon (nomLeu1), macaque (rheMac3), and baboon (papHam1). We used the UCSC genome browser’s liftOver program to align each ExAC SNP along with its 60bp sequence context to the six aforementioned reference genomes. We used the baboon reference genome solely for the ascertainment of all other substituted-species categories (rather than including a substituted-baboon category in the analysis).

For gene annotations, we downloaded the refGene table of the RefSeq Genes track from the UCSC genome browser. For each SNP in our data, we extracted all gene isoforms in which the position was included. We kept all ExAC SNPs that fell in a coding exon, intron or untranslated region. We excluded from the analysis non-autosomal SNPs, SNPs that had multiple annotations corresponding to different transcript models, and SNPs with a sample size of less than 100,000 chromosomes. After applying the filters we were left with 6,002,065 SNPs.

For recombination rates, we downloaded the sex-averaged recombination rate map constructed by Kong et al. [87], which estimates rates at a resolution of 10kbp bins.

### The probability of ancestrally shared polymorphisms

In order to construct an upper bound on the probability of a human polymorphic site being ancestrally shared with another species, we consider the case of a selectively neutral polymorphism shared with chimpanzees. A polymorphism observed in the human sample at the current time is an ancestral polymorphism at the time of the human-chimpanzee split only if there are at least two lineages ancestral to the human sample at the human-chimpanzee split time. Leffler et al. [42] assume a constant human effective population size of 10,000 people throughout history, and estimate a probability of about 1.0 · 10^−5^. In the **Supplementary Material**, we augment Leffler et al.’s approximation with more complex demographic models for recent human history and derive an upper bound of 1.4 · 10^−5^ for this probability. Multiplying this probability by the number of exonic sites (3,531,936) in our data, we get an expected number of 49 sites in our data that are ancestrally shared with chimpanzees.

However, our derivations are based on a pre-out-of-Africa effective population size 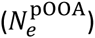 of 10,000 people. Very little is known about human demographic history prior to the out-of-Africa event, and as we show in the **Supplementary Material**, the probability of an ancestral polymorphism rises very quickly with increasing 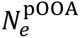. Estimates of 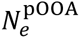 range between 7300 [88] and 12,500 [89] people and are continually revised as estimates of human mutation rate and demographic history are refined. With 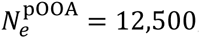, we get an upper bound of 283 polymorphisms in the dataset that are expected to be shared between human and chimpanzee, which compose at most 1.2% of the substituted-chimpanzee sites. The upper bound for other species, or sites under purifying selection, should be even smaller, and are overall too few to affect our results. We therefore conclude that ancestral polymorphisms are too few to significantly affect our analysis.

### Simulating mutational models and demographic models

To get a theoretical expectation for the fraction of rare variants under different mutational models, we used various software for computing the expected sample SFS of 33,750 diploid individuals, corresponding to the size of the non-Finnish European subsample in the ExAC dataset. For all mutational models, which we describe below, we generated predictions under various demographic models from recent literature: Gazave et al. [90] (model 2 in their work), Tennessen et al. [16] and Nelson et al. [14].

For the infinite-sites model, we computed the expected sample SFS analytically using `fastNeutrino` [66]. The infinite-sites model corresponds to an upper bound for the fraction of rare variants, but nonetheless predicted a fraction of rare variants much lower than that observed in data (75%-78%) for all non-CpG mutations under the Gazave et al. (59%) and Tennessen et al. (60%) demographies. The Nelson et al. model, which was inferred using a larger sample size of 11,000 people predicted 75% of biallelic polymorphisms would be rare under the infinite-sites model. In order to fit the highest observed fraction of rare variants for non-CpG sites in the ExAC data, we modified the parameters of the most recent epoch of exponential growth in Nelson et al. We estimated these parameters using `fastNeutrino` [66] on all A->C intronic mutations from ExAC. The inferred parameters were: current effective population size of 4,009,877 diploids, and an exponential growth onset time of 119.47 generations in the past with a growth rate of 5.38% per generation. The more ancient demographic parameters were fixed to the same values as in the model of Nelson et al.

We assume multiple mergers (non-Kingman merger events) have negligible effect on the SFS since the sample size is significantly smaller than the current effective population size. A similar demographic model [91] with a four-fold smaller current effective population size exhibited a relative difference of only about 1.3% and 0.3% in the proportion of singletons and doubletons respectively for a comparable sample size of 50,000 people. Hence, we felt confident in using the Kingman coalescent for drawing genealogies.

For the finite sites model, we first simulated independent coalescent trees using `ms` [92] and then generated 1kb non-recombining sequences for each coalescent tree using the desired recurrent mutation rate with the 4 x 4 Jukes-Cantor model of mutation [60]. We used the program `Seq-Gen` [93] to drop recurrent mutations on coalescent trees drawn from `ms`. We used mutation rates in a uniform logarithmically spaced grid of 40 points ranging from 10^−9^ to 5.3 · 10^−5^ mutations per basepair per generation per haploid. For each value of the mutation rate, we simulated enough sequence data so that at least 100,000 biallelic polymorphic sites were available to reliably estimate the expected fraction of rare variants. If we indicate whether a variant is rare by *Y*, then for each mutation rate *μ*, the expected fraction of rare variants is

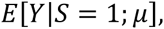

where *S*, is an indicator variable indicating whether a site is polymorphic and, specifically, biallelic. Finally, we considered a model with additional, within-mutation-type heterogeneity in mutation rate. Specifically, we considered a model in which sites of a particular mutation type (e.g., C->A sites) have a mean mutation rate *μ* as before, but the mutation rate itself, *M*, is no longer fixed (and equal to *μ*), but rather a random variable with mean *μ*. Let *f*(*M*|*S* = 1; *μ*) be the probability density function of *M* in a site with mean mutation rate *μ* conditional on it being biallelic. Then, by the law of total expectation we have:

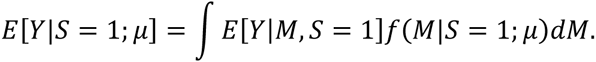

By Bayes’ rule, *f*(*M*|*S* = 1; *μ*) is determined by both the within-mutation-type distribution of mutation rates, *g*(*M*; *μ*), and the probability of a site with mutation rate *M* being biallelic, as follows:

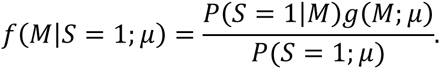

Therefore,

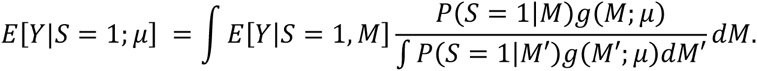

For a large range of *M*, we have already estimated *E*[*Y*|*M*] as described above. From the same simulations we have estimated the probability of a site with mutation rate *M* being a biallelic polymorphism, *P*(*S* = 1|*M*). Lastly, the distribution of mutation rates due to within-mutation-type variance was modeled using a lognormal distribution:

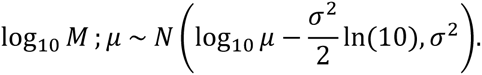

The mean parameter in the lognormal distribution above ensures that *E*[*M*] = *μ*. *σ* was arbitrarily chosen to be 0.57 (red line in Figure 4A). Notably, Hodgkinson et al. also fit a lognormal distribution of mutation rates to their dataset of co-occurrence of SNPs in chimpanzees and humans, and estimate a similar value of σ = 0.83 for non-CpG mutations [64] and *σ* = 0.8 for CpG transitions (personal correspondence).

### Logistic model for the probability of a variant to be rare

We tested whether the species trend across cSFS is due solely to the effect of mutation rate variation. We used a logistic regression model to examine whether a residual substituted-species trend remains after controlling for mutation type. Let be a binary-valued random variable indicating whether a variant is rare, 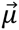 be a vector of mutually exclusive indicator (dummy) variables for each mutation type, 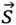 be a vector of mutually exclusive indicator variables for the divergence pattern for the variant (substituted in one of the primates or human-private) and *Z* be an indicator of whether the variant is nonsynonymous (we only considered coding variants). We fitted the logistic regression model

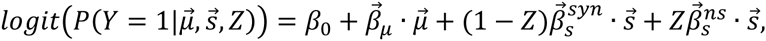

where the parameters 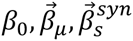, and 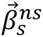 were learned from the data. We tested whether the coefficients 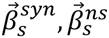 exhibit a trend across *s*, i.e. whether the probability of the variant being rare is associated with the relatedness of the substituted species. When ignoring the mutation rate effect (i.e. fixing 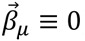 the 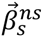 estimates were perfectly anti-correlated with the relatedness of the substituted species to human, consistent with the observation in data (Figure 6A). We then allowed for an effect for the mutation type by estimating 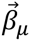 for the different categories of mononucleotide mutation types (Figure 6B). We also estimated a model with a finer resolution of mutational categories, further partitioning the mononucleotide mutation types by their two flanking nucleotides (Figure 6C). For nonsynonymous sites, which likely involve the strongest purifying selection pressures, the trend persisted even after controlling for mutation rate variation (Spearman *ρ* = 1, *p* = 0.016 for both mononucleotide correction and for the correction including flanking nucleotides context).

## Acknowledgements

We thank ExAC and the groups that provided exome variant data for comparison. A full list of contributing groups can be found at http://exac.broadinstitute.org/about. We also thank Molly Przeworski, Ziyue Gao, Doc Edge, Xun Lan, David Golan, Kelley Harris, Anil Raj and Eyal Elyashiv for helpful comments on the manuscript and/or valuable discussions.

